# Assessment of single cell RNA-seq normalization methods

**DOI:** 10.1101/064329

**Authors:** Bo Ding, Lina Zheng, Wei Wang

## Abstract

We have assessed the performance of seven normalization methods for single cell RNA-seq using data generated from dilution of RNA samples. Our analyses showed that methods considering spike-in ERCC RNA molecules significantly outperformed those not considering ERCCs. This work provides a guidance of selecting normalization methods to remove technical noise in single cell RNA-seq data.

## Introduction

Single cell RNA-seq (scRNA-seq) is becoming a powerful tool to study many biological problems such as tissue heterogeneity(1– 3) and cell differentiation during development(4– 6,7). Normalization is particularly critical for interpreting scRNA-seq data including detection of differentially expressed gene and identification of cell sub-types(7). Small amount of samples used in scRNA-seq often leads to higher technical noise compared to bulk RNA-seq, which needs to be removed. Various methods have been developed for normalization of scRNA-seq data including fragments per kilobase of transcript per million mapped reads (FPKM)(8), upper quartile (UQ)(9), Trimmed mean of M-values (TMM)(10), DESeq(11), removed unwanted variation (RUV)(12), and gamma regression model (GRM)(13). Most of them were developed for bulk RNA-seq and then directly applied to scRNA-seq analysis. The performance of these methods was assessed on bulk(12, 14) but not on scRNA-seq data because the ground truth is likely unknown for single cell assays.

Recently, the NIH Single Cell Analysis Program - Transcriptome Project (SCAP-T) collected two well-characterized human reference RNA samples: Universal Human Reference RNA (UHR) and Human Brain Reference (HBR)(15). A set of RNA-seq data were generated using different amount of RNAs obtained from dilution (10ng considered as bulk, 100pg and 10pg). These samples were prepared for sequencing using three protocols: antisense RNA IVT protocol (abbreviated as aRNA or A)(16), a customized C1 SMARTer protocol performed on a Fluidigm C1 94-well chip (S)(17), and a modified NuGen Ovation RNA sequencing protocol (N)(18) (Table 1). These data can be divided into six groups based on the sample source and amount: UHR, bulk; UHR, 100pg; UHR, 10pg; HBR, bulk; HBR, 100pg and HBR 10pg. It is natural to assume that 100pg samples would be more similar to bulk than 10pg samples. Therefore, this set of data is invaluable to compare normalization methods because the ground truth is known. Furthermore, 51 of these samples were mixed with spike-in ERCC RNA molecules, which makes it possible to evaluate the impact of considering ERCCs in normalization of scRNA-seq data.

**Table 1.** Dilution RNA-seq data generated by SCAP-T.

| Single Cell Protocol | Human brain reference RNA amount (HBR) |  |  | Universal human reference RNA amount (UHR) |  |  | Total |
| --- | --- | --- | --- | --- | --- | --- | --- |
|  | 10pg | 100pg | bulk | 10pg | 100pg | bulk |  |
| aRNA (A) | 18(5)* | 3 | - | 12 | 7 | - | 40 |
| C1 SMARTer (S) | 15(15) | 5(5) | - | 15(15) | 5(5) | - | 40 |
| Nugen Ovation RNASeq V2 (N) | 4 | 4 | - | 15 | 11 | - | 34 |
| None | - | - | 3 | - | - | 3 | 6 |
| Total | 37 | 12 | 3 | 42 | 23 | 3 | 120 |

\*The number in the parenthesis is the number of samples containing spike-in ERCCs.

## Results

We selected seven methods to normalize the data of the 120 RNA-seq samples. These methods fall into two categories, consideration and not consideration of spike-in ERCC molecules in normalization.

### Evaluation of the methods not considering ERCC

We first evaluated the performance of five methods not considering ERCCs, FPKM(8), UQ(9), TMM(10), DESeq(11), RUVr(12) (non-considering ERCC version of RUV) (see Methods for the details of setup for running each method). 13,375 genes were selected with log(fpkm)>2 in at least 2 of the 114 non-bulk samples (100pg or 10pg). Using Pearson correlation as the similarity metric, we clustered all the 120 samples (114 non-bulk and 6 bulk samples) using hierarchical clustering (Figure 1, Figure S1). For all the methods, the UHR and HBR samples are well separated as expected and we thus focused on evaluating how well the HBR and UHR samples are further clustered. In either the UHR or HBR group, the normalized samples by FPKM, UQ, TMM and DESeq tend to cluster by sequencing protocols rather than RNA amount, which indicates the difference introduced by the protocols were not completely removed by normalization. In contrast, the HBR samples normalized by RUVr were largely clustered by RNA amount rather than sequencing protocol, although a significant portion of 10pg samples sequenced using aRNA were clustered separately (Figure 1 and S1). Qualitatively, RUVr normalization alleviates difference between scRNA-seq protocols and the clustering results are closer to the ground truth than the other methods, i.e. the samples are clustered based on the source (HBR) and the RNA amount (bulk, 100pg or 10pg). However, the UHR samples normalized with RUVr were still clustered according to protocols rather than RNA amount. The other four methods showed worse clustering results than RUVr because 10pg and 100pg samples were mixed in each protocol group (Figure 1 and S1).

**Figure 1.**
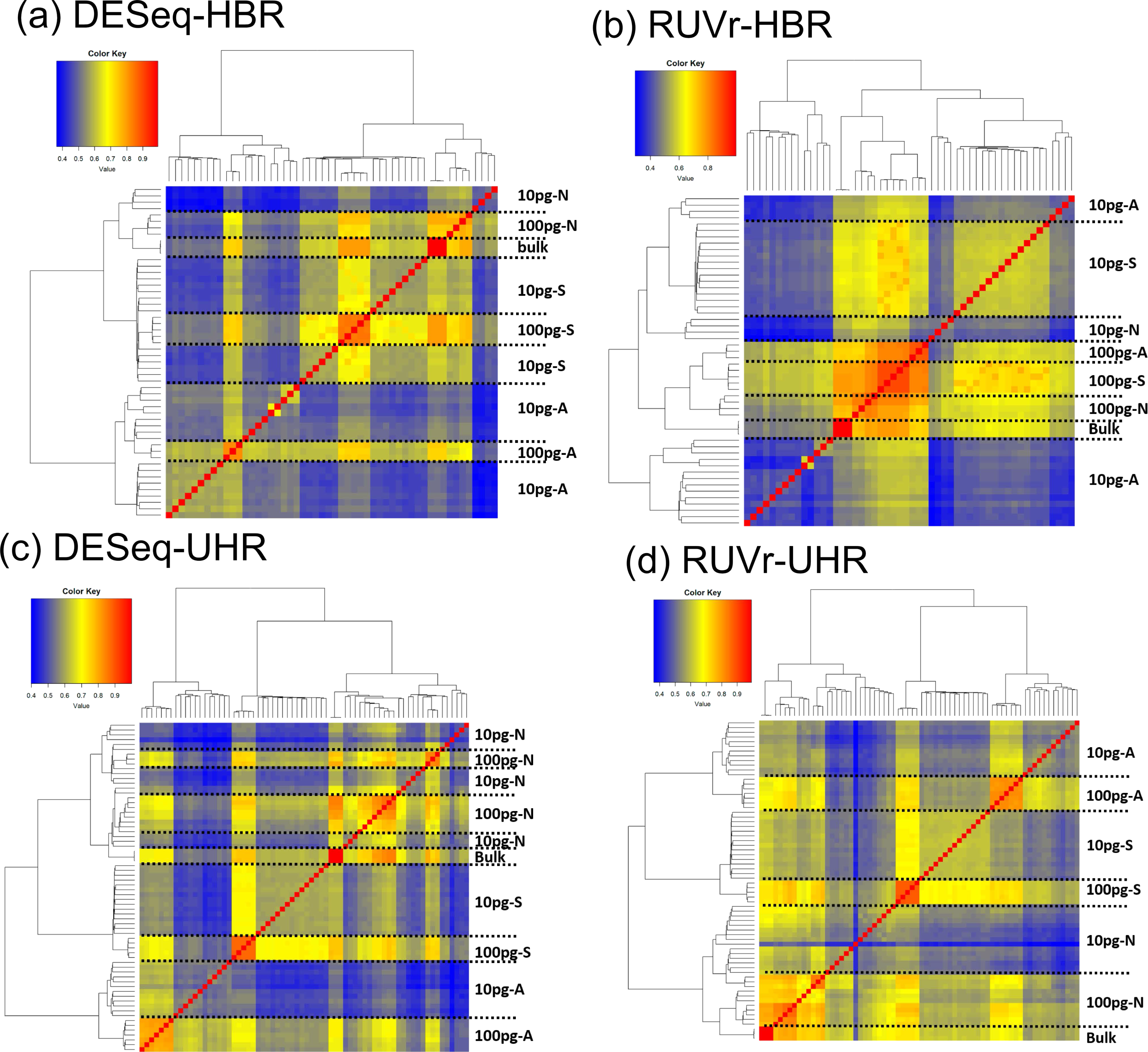
Hierarchical clustering of HBR and UHR RNA-seq samples with different normalization methods not considering ERCC. (a) DESeq with HBR samples; (b) RUVr with HBR samples (c-d) DESeq and RUVr with UHR samples. The results of other methods are shown in Figure S1. A, aRNA; N, Nugen Ovation RNASeq V2; S, C1 SMARTer.

To quantify the similarity between the clusters generated from different normalization methods and the ground truth, we cut the clusters at different hierarchy levels and calculated the Rand index between the clusters and the ground truth, which reflects the correctness of clustering. Rand index is the ratio between the sample pairs that are correctly clustered and the total sample pairs(19). Because all the methods successfully separated HBR and UHR samples as discussed above, we calculated Rand index on HBR and UHR samples separately. For either the HBR or UHR samples, we varied the cutoffs to cut the hierarchical trees into 2 to 7 clusters and calculated Rand index for each cutoff (Figure 2(a), Figure S2). Obviously, RUVr outperformed the other methods.

**Figure 2.**
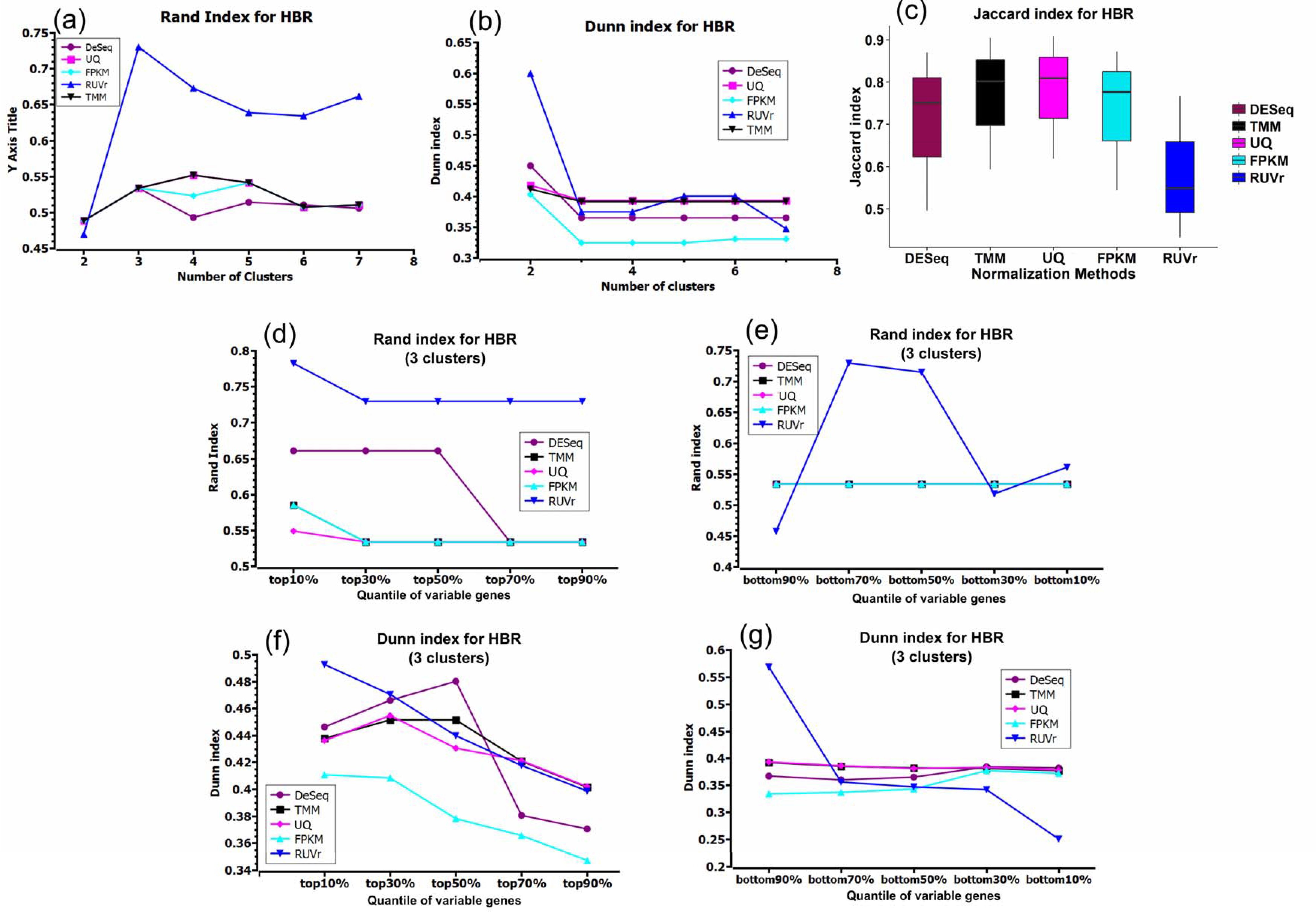
Comparison of 5 normalization methods not using ERCC on the HBR samples. (a) Rand index with different number of clusters; (b) Dunn index with different number of clusters; (c) Jaccard index with three clusters; (d) Rand index with the most variable genes; (e) Rand index with the least variable genes; (f) Dunn index with the most variable genes; (g) Dunn index with the least variable genes. (d)-(g) are results using 3 clusters as the ground truth (bulk, 100pg and 10pg HBR samples). UHR results are shownin the Supplementary Materials (Figure S2).

Next, we evaluated the compactness and separation of clusters using Dunn index. Dunn index is the ratio of the smallest distance between clusters to the largest distance of intra-cluster, and the distance between the twosamples is defined as 1 minus Pearson correlation between the samples(20) We computed the Dunn index on HBR and UHR samples separately. RUVr had a much higher Dunn index than the other methods while all the samples were clustered into two groups. For more clusters, all the methods had comparable Dunn index. This observation suggests that the separation between clusters is insensitive to normalization methods when the 120 samples were clustered into more than 3 clusters.

It is important to examine whether the clustering results are robust tothe selection of samples and genes. We first divided HBR or UHR samples into three clusters: bulk, 100pg and 10pg. Then we randomly selected 50% of the total samples, clustered them into three groups using the 13,375 genes selected above. We computed the maximum Jaccard index(21) (see definition in Methods) between the new cluster and the ground truth. The Jaccard index reflects the correctly clustered samples using the subsets of samples and thus the robustness of the clustering. All methods but RUVr showed comparable Jaccard index. Next, to evaluate the robustness of clustering to genes used to calculate Pearson correlation coefficient between samples, we ranked the 13,375 genes using their coefficients of variant (CV)(22), which is the standard deviation of gene expression divided by the mean, reflecting the variation of a gene's expression across samples. We selected 10 sets of genes: top 10%, top 30%, top 50%, top 70%, top 90% and bottom 10%, bottom 30%, bottom 50%, bottom 70%, and bottom 90%. In general, using the most variable genes achieved the most correct clustering, i.e. high Rand index, and using a sufficient number of most variable genes (top 10% for RUVr and FPKM, top 30% for UQ and TMM, top 50% for DeSeq) gave the highest Dunn index that represents the best separation of the clusters (Figure 2(d)-(g)). Using the least variable genes (most invariable genes) such as the bottom 10% variable genes normally showed lower Rand and Dunn index than using more variable genes such as the bottom 90% variable genes (Figure 2(d)-(g)).

### Evaluation of the methods considering ERCC

We next evaluated the performance of two methods considering ERCCs: RUVg (RUV model considering ERCC) (12) and GRM(13) (see Methods for the details of the setup for running each method). Among the 120 RNA-seq runs, 45 samples containing spike-in ERCCs were normalized using the two methods and then clustered with 6 bulk samples together (Figure 3). The clustering results were similar to the ones using the 13,375 selected genes. UHR and HBR samples were clearly separated and the samples with the same amount of RNA were largely clustered together. One outlier in the RUVg cluster is a HBR 10pg sample sequenced using the aRNA protocol that was clustered with HBR 100pg samples.

**Figure 3.**
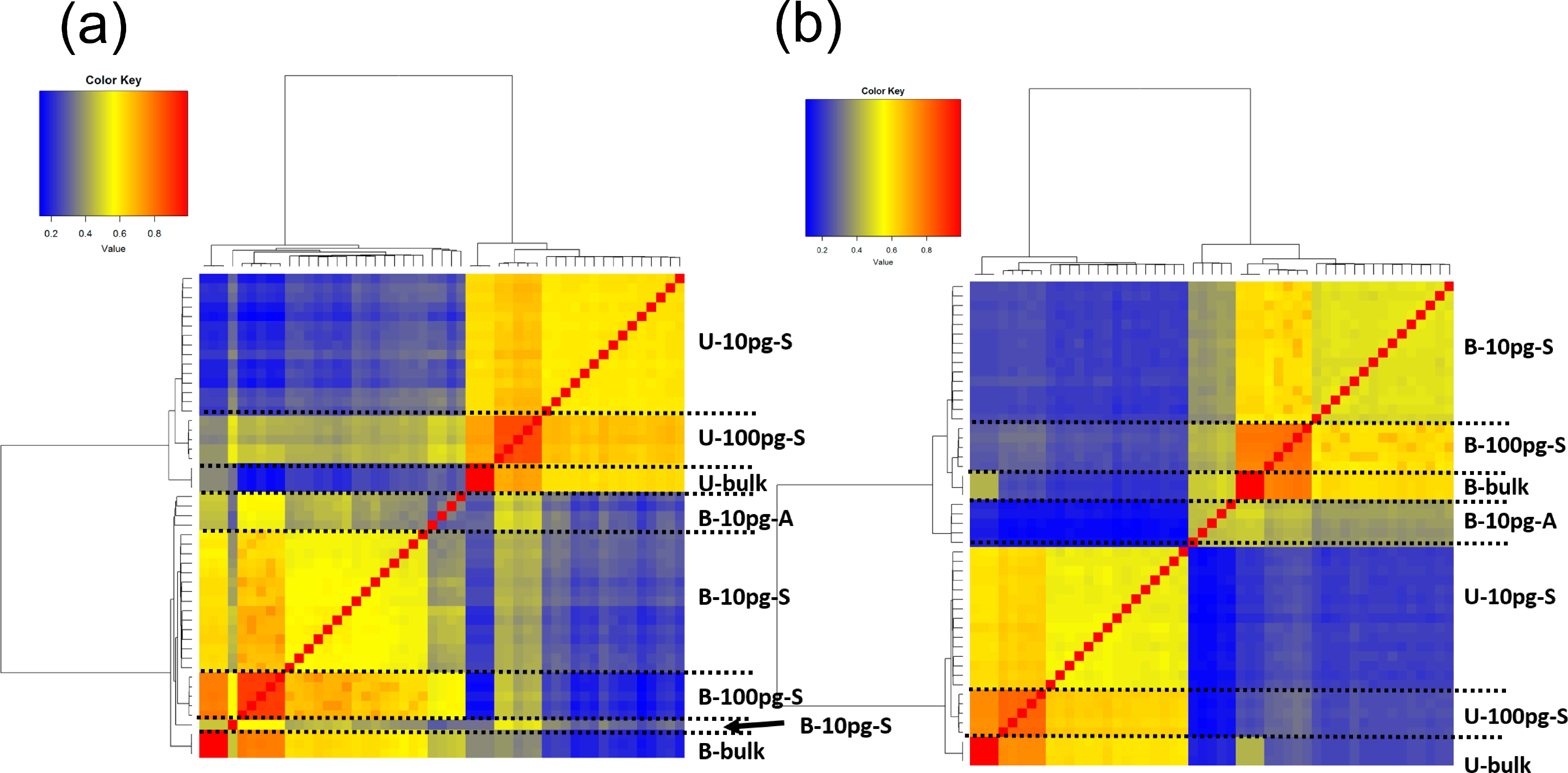
Hierarchical clustering of 51 samples with different normalization methods using ERCC. (a) RUVg; (b) GRM. B, HBR; U, UHR; A, aRNA; S, C1SMARTer.

GRM showed slightly higher Rand index than RUVg when the number of cluster is same as the ground truth (6) or more (Figure 4(a)). GRM also had better Dunn index and comparable Jaccard index (Figure 4(b) and (c)), which suggests that GRM achieves better separation of clusters (Dunn index). The robustness to the sample selection is comparable for the two methods as indicated by their similar Jaccard index. Furthermore, we assessed how sensitive the two methods are to gene selection. For Dunn index, GRM outperformed RUVg regardless whether the most variable or the least variable genes were selected (Figure 4(d) and (e)). For Rand index, using the most variable genes, GRM consistently showed higher values than RUVg; using the least variable genes, GRM achieved higher Rand index using bottom 90% and 70% genes but lower values using the bottom 50%, 30% or 10% genes than RUVg and RUVr (6 clusters of bulk, 100pg and 10pg samples for HBR and UHR were considered as the ground truth) (Figure 4(f) and (g)). We examined the clusters generated using the bottom 30% variable genes. GRM still correctly clustered all the samples but RUVg incorrectly clustered some HBR samples with UHR despite of a higher Rand index (Figure 5). We then used the bottom 10% variable genes (the most invariable genes) for a further comparison. Both methods mis-clustered UHR bulk with the HBR samples but RUVg had additional HBR samples mistakenly clustered with UHR samples (Figure 5). For Jaccard index, GRM performed better than RUVg and RUVr with genes selected from top 10% through top 90% and bottom 90%, bottom 70% and bottom 50%. When the least variable genes were selected (bottom 10%), RUVg and RUVr significantly outperformed GRM. (Figure S3).

**Figure 4.**
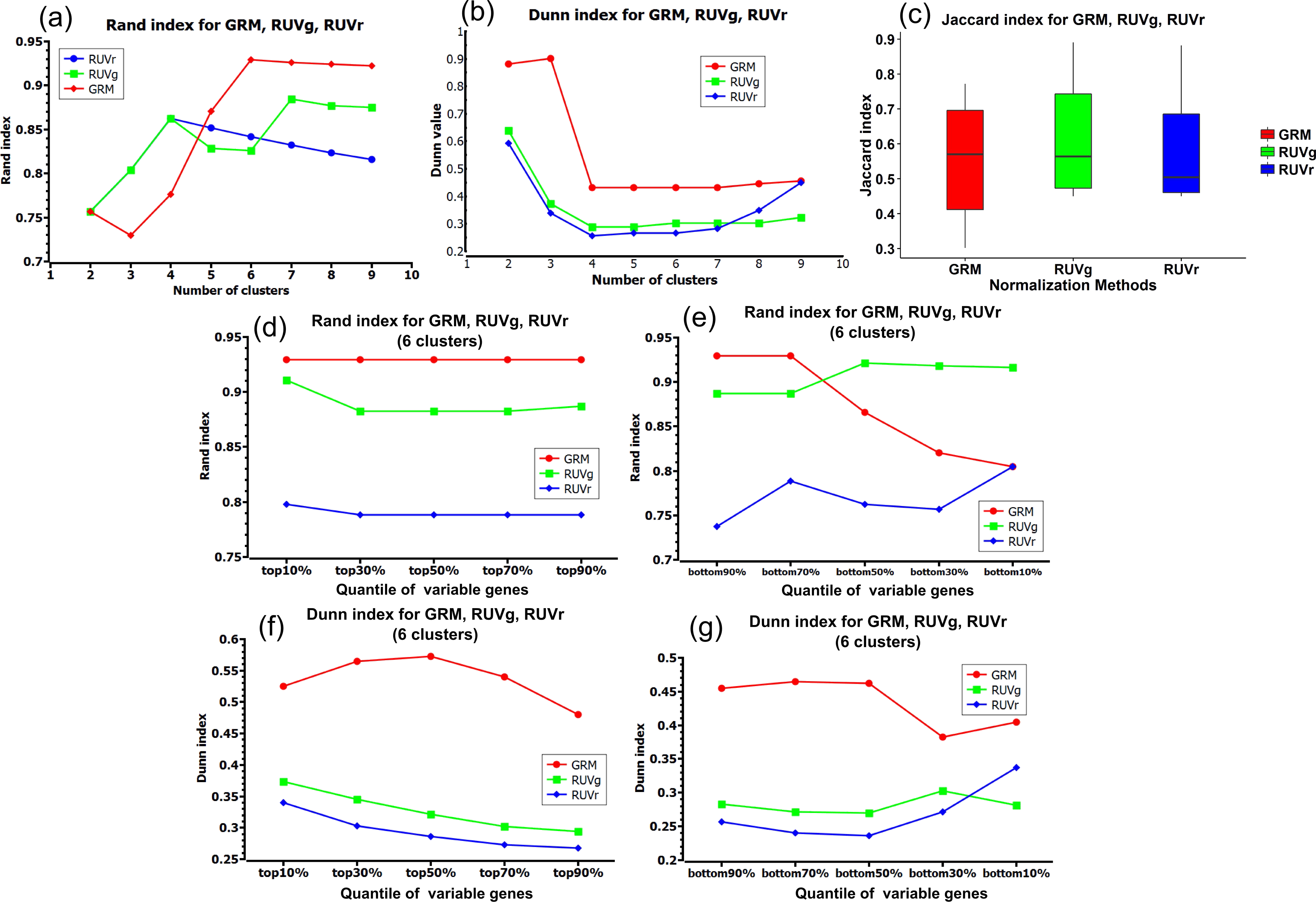
Comparison of two normalization methods using ERCC (RUVg and GRM) and one not using ERCC (RUVr). (a) Rand index with different number of clusters; (b) Dunn index with different number of clusters; (c) Jaccard index with six clusters; (d) Rand index with the most variable genes; (e) Rand index with the least variable genes;(f) Dunn index with the most variable genes; (g) Dunn index with the least variable genes. (d)-(g) are results of using 6 clusters as the ground truth (bulk, 100pg and 10pg of HBR and UHR samples).

**Figure 5.**
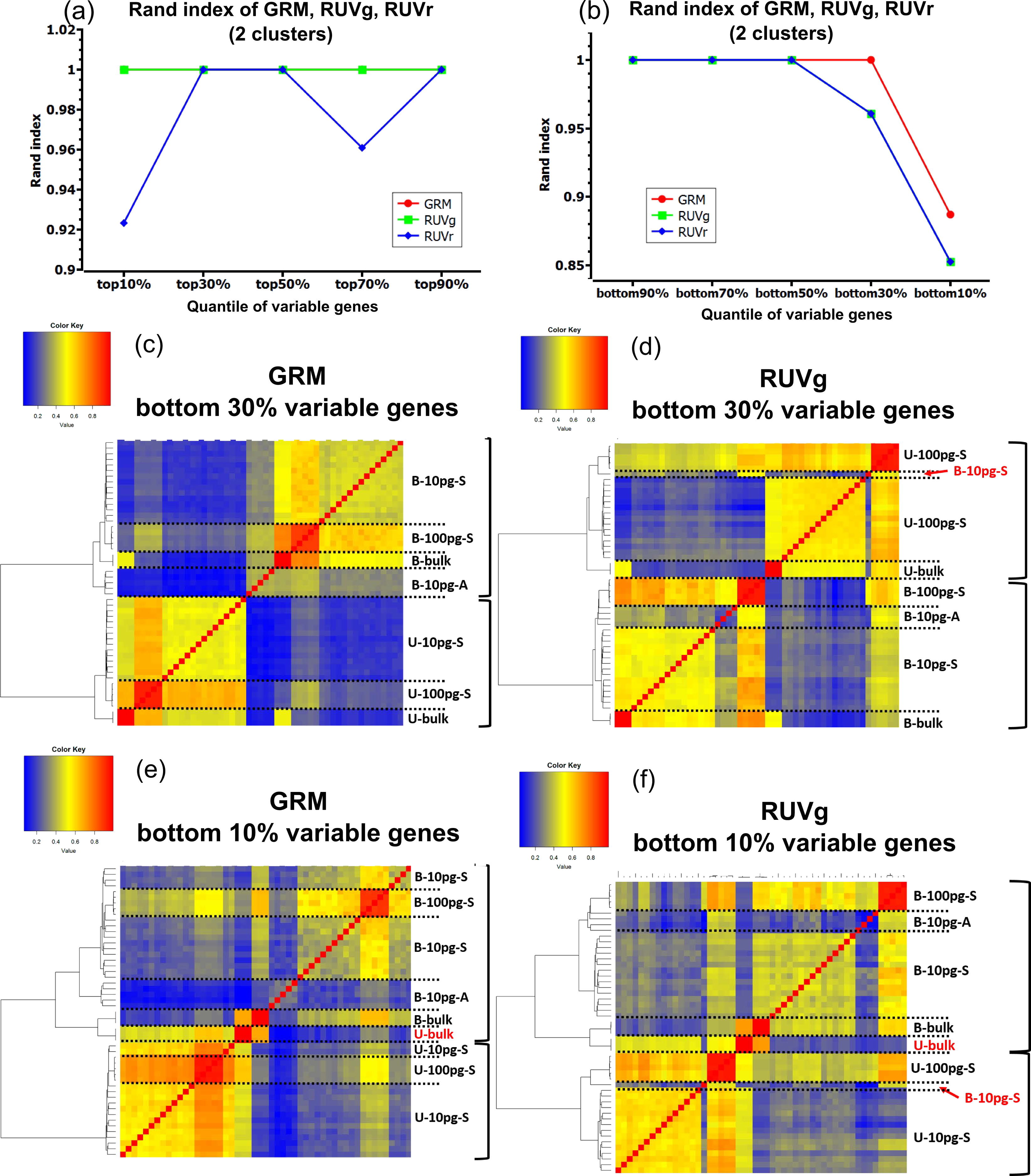
Comparison of two normalization methods using ERCC (RUVg and GRM) and one not using ERCC (RUVr) if the ground truth has two clusters (HBR and UHR samples). (a) Rand index of GRM, RUVg and RUVr with the most variable genes. (b) Rand index of GRM, RUVg and RUVr with the least variable genes. (c)-(d) hierarchical clustering of GRM and RUVg using the bottom 30% variable genes. (e)-(f) hierarchical clustering of GRM and RUVg using the bottom 10% variable (the most invariable) genes. Right brackets label the first branches. Red samples are wrongly clustered samples.

Notably, both methods considering ERCC outperformed the methods not considering ERCC on the 51 samples (45 10pg or 100pg and 6 bulk samples) containing spike-in ERCCs (Figure 4). We took RUVr as a representative method and it largely outperforms the other methods not considering ERCCs. Obviously, both GRM and RUVg showed higher Rand, Dunn and Jaccard index than RUVr, which indicates the value of considering ERCC in normalization and denoise of scRNA-seq.

## Discussion

Here, we evaluated the performance of seven normalization methods on 120 RNA-seq runs in terms of correctness (Dunn index), compactness (Rand index) and robustness (Jaccard index, robustness analysis using different sets of samples and genes) of clustering. The results showed that, for methods not considering spike-in ERCCs, RUVr showed higher Dunn index and lower Jaccard index than FPKM, UQ, DESeq and TMM; all methods showed similar Rand index. Considering ERCC such as in the models of RUVg and GRM significantly improved the performance. Between RUVg and GRM, GRM is more robust in terms of selecting different sets of genes that generate similar clusters.Spike-in ERCCs would reduce the sequencing depth of mRNAs of interest and there is also a concern about whether the synthesized ERCC molecules behave same as the mRNAs from the cell. Our analyses suggest that calibration of scRNA-seq to the spike-in ERCC is powerful to remove technical noise when the ERCCs are correctly modeled.

## Methods

### Selection of variable genes

Before applying different normalization methods on the data, we first selected variable genes using the following criteria: 1) across the 114 non-bulk samples, at least 2 samples with log(fpkm)>2; 2) across 114 non-bulk samples, variant of log(fpkm) larger than 1. This way, we found 13,375 genes for the following analyses.

### Running normalization methods

Fragments per kilobase of transcript per million mapped reads (FPKM)(23): Fragments per kilobase of transcript per million mapped reads. FPKM normalizes the gene counts in consideration of library size and gene length. Upper Quartile (UQ)(9) and Trimmed mean of M-values (TMM)(10) methods are implemented in edgeR Bioconductor package, and we run the R function using default parameters. The DESeq(24) method is implemented in the Bioconductor package. We run the R function using default parameters. Remove unwanted variants (RUV)(25) is included in the RUVnormalize Bioconductor packages. The model sets up a generalized linear regression model between observed RNA-seq read counts and the known covariates of interest along with unknown unwanted variation factors. Different options of RUV, RUVr and RUVg, have different ways to estimate the unwanted variant factors. RUVr uses residuals from a first-pass regression of read counts(25) and considers the lest differentially expressed genes across samples. RUVg considers negative controls such as ERCC spike-ins and assumes that they are not differentially expressed across samples. All the R functions were run using default parameters. For empirical undifferentially expressed genes in running RUVr, we chose genes not in the top 6000 differentially expressed genes. GRM(26) fits a gamma regression model between FPKM values of reads and the concentration of ERCC spike-ins, and then make estimates of the molecular concentration of the genes from the reads.

### Assessment of clustering performance

After normalization, we clustered the normalized gene reads using hierarchical clustering with the Ward method and the metric was Pearson correlation between normalized gene reads. We assessed the clustering results using the following statistical indices:

1. Rand index(19) evaluates the correctness of clustering using prior labels. Given a set of n elements S={*s*_1_,*s*_2_, …,*s_n_*} and two partitions needed to be compared, a partition of S into M clusters *X*={*X*_1_,*X*_2_,…,*X_M_*} and a partition of S into N clusters, for certain *Y* = {*Y*_1_,*Y*_2_,…,*Y_N_*} for certain 1 ≤ *_i,j_* ≤*n*(*i* ≠ *j*),1 ≤ *k,k_1_,k_2_* ≤ *M*(*k*_1_ ≠ *k*_2_), 1≤ *l,l_1_,l_2_* ≤ *N*(*l*_1_ ≠ *l*_2_), *a*, *b*, *c*, *d* are defined as following:

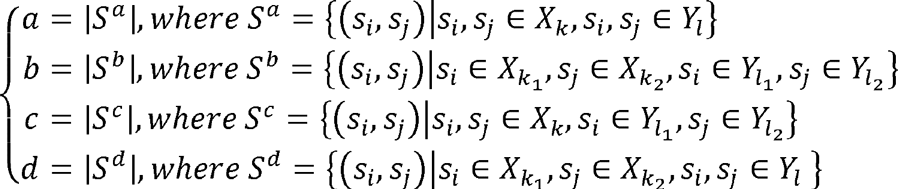 Then Rand index can be computed as following:

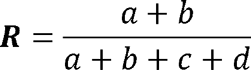

where (a+b) is the number of agreements between X and Y, while (c+d) is the number of disagreements between X and Y. The higher Rand index is, the more similar the two partitions are. As we compared the clustering partitions to the prior known labels, the higher Rand index indicates the better the clustering is.
2. Dunn index(27) evaluates the compactness and separation of the clustering. Specifically, given a set of n elements S={*s*_1_,*s*_2_,…,*s_n_*} and the clustering partition as ζ = {*C*_1_,*C*_2_,…*C_k_* Dunn index is computed as following:

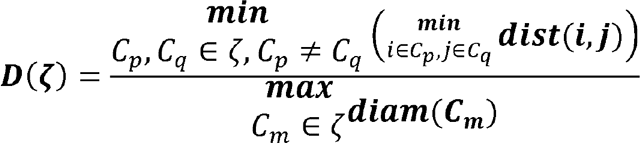

By definition, the higher the Dunn index is, the better the clustering is, considering enough separation compared to diameter of individual clusters. Here, the distance between two samples are defined as 1 minus Pearson correlation coefficient between two samples.
3. Jaccard index(28, 29) evaluates the stability of clustering that measures the similarity between two finite subsets. Given two subsets A and B, the Jaccard index is computed as:

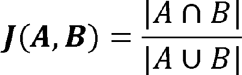

We used the Flexible Procedures for Clustering (fpc) package in R to calculate the Jaccard index with randomly subsetting without replacement of samples. For each time drawing a subset of samples, we clustered the samples and then computed the maximum Jaccard index between the new cluster and the prior known cluster. Repeating B times for drawing B random subsets (here we chose B=100), we then computed the mean of all the maximum Jaccard index.(30) The higher the Jaccard index is, the more robust the clustering is.

## Acknowledgement

We are grateful to the SCAP-T Consortium to provide access to the dilution data. This work is partially supported by NIH (U01MH098977).

